# Differential effects of diphenyl diselenide (PhSe)_2_ on mitochondria-related pathways depending on the cellular energy status in Bovine Vascular Endothelial Cells

**DOI:** 10.1101/2024.06.14.599060

**Authors:** Letícia Selinger Galant, Laura Doblado, Rafael Radi, João Batista Teixeira da Rocha, Andreza Fabro de Bem, Maria Monsalve

## Abstract

Cellular energy metabolism varies depending on tissue and cell type, as well as the availability of energy substrates and energy demands. We recently investigated the variations in cellular metabolism and antioxidant responses in primary bovine vascular endothelial cells (BAECs) under different energetic substrate conditions *in vitro*, specifically glucose or galactose. In this context, pharmacological agents may affect cells differently depending on their energy metabolism status. In this study, we aimed to characterize the effects of diphenyl diselenide ((PhSe)_2_), a redox-active molecule known for its prominent cardiovascular effects, on redox-bioenergetic cellular pathways under glycolytic or oxidative conditions in BAECs. Under glucose conditions, (PhSe)_2_ positively impacted mitochondrial oxidative capacity, as assessed by respirometry, and was associated with changes in mitochondrial cellular dynamics. However, these changes were not observed in cells cultured with galactose. Although (PhSe)_2_ induced the nuclear translocation of the redox sensitive nuclear factor erythroid 2-related factor 2 (Nrf2) in both glucose and galactose media, Nrf2 remained in the nuclei of cells cultured in galactose for a longer duration. Additionally, activation of another redox sensitive transcription factor, forkhead O3 (FOXO3a) was only detected in galactose media. Notably, (PhSe)_2_ induced the expression of genes controlling mitochondrial antioxidant capacity and glutathione synthesis and recycling in glucose media, whereas its effects in galactose media were primarily focused on glutathione homeostasis. In conclusion, our findings underscore the critical influence of cellular metabolic status on the antioxidant capacity of redox-active molecules such as (PhSe)_2_.

## 1. Introduction

Cellular energy metabolism may vary depending on the tissue and cell type, the energy substrates available, and the energy demands. Cancer cells, in particular, commonly exhibit a more avid non-oxidative glycolytic metabolism than non-tumor cells [1]. However, they also need to adapt their metabolism in response to substrate availability, which results in corresponding changes in cell physiology and redox-bioenergetic status [2]. Similarly, pharmacology agents may have different cellular effects, depending on the energy metabolism status of the target cells [3].

The use of different cell culture media conditions provides a suitable model for evaluating cellular metabolic performance *in vitro*. While both glucose and galactose are good cellular energy sources, galactose enters the glycolytic pathway through the energy-consuming Leloir pathway [4]. When glucose in the culture medium is replaced by galactose (5-10 mM), cells are forced to shift to the mitochondrial oxidative metabolism to meet their energy demands. This condition not only activates mitochondrial oxidative phosphorylation OXPHOS but fully reprograms mitochondrial function and dynamics and facilitates the evaluation of mitochondrial susceptibility to various agents, revealing sensitivities related to oxidative metabolism that are obscured when cells are cultured in high glucose-containing medium [5,6]. In contrast, glucose catabolism tends to be primarily non-oxidative when glucose levels reach a certain threshold, known as the “Crabtree effect.” As a result, cells grown in media with a standard of glucose concentrations (25 mM) tend to acquire a highly non-oxidative glycolytic phenotype [7].

The activation of the mitochondrial OXPHOS is controlled by the master transcriptional coactivator peroxisome proliferator-activated receptor gamma coactivator -1alpha (PGC-1α) and linked to the reprogramming of the cellular capacity to regulate its REDOX status [8]. Furthermore, Nrf2 and FOXO3, transcription factors that control antioxidant gene expression, and respond to oxidative stress translocating to the nucleus [9,10], are also well known to be sensitive to the cellular metabolic status [11,12], and are regulated by PGC-1α [13,14]. However, studies that evaluate the redox activity of molecules rarely consider the differential effect of the cellular metabolic status. In a recent study, we observed that endothelial cells maintained in a galactose-containing medium, not only exhibited higher mitochondrial oxidative capacity and intercellular metabolic coupling than were cultured in galactose but also showed differences in the nuclear levels of Nrf2 and FOXO3 which may be indicative of differential sensitivity to molecules with redox activity [15], highlighting the relevance of redox-metabolic coupling and the need to control for metabolism in cell assays. Accordingly, it is reasonable to expect that pharmacological drug protocols will have different cellular effects depending on the energy metabolism status of the target cells.

(PhSe)_2_ is a promising organoselenium compound with antiatherogenic properties [16–18]. The cardiovascular actions of (PhSe)_2_ have been previously tested in several *in vivo* and *in vitro* models, including hypercholesterolemic mice and rabbits, human low-density lipoprotein (LDL), and vascular endothelial and macrophage cells [19,20]. (PhSe)_2_, mechanism of action has been proposed to be related to its antioxidant activity. It has been shown to react with reactive cellular redox-sensitive thiols, such as cysteine and the tripeptide glutathione (GSH), an important antioxidant, functioning as a glutathione peroxidase (GPx) mimic, and facilitating the recycling of reduced glutathione. Additionally, (PhSe)_2_ promotes the oxidation of critical cysteinyl residues in kelch-like ECH-associated protein 1 (Keap1), a redox-sensing protein that retains Nrf2 in the cytoplasm. This oxidation leads to the translocation of Nrf2 into the nucleus and its subsequent activation [16,21–23]. Furthermore, (PhSe)_2_ has been shown to increase the levels of two key mitochondrial antioxidant enzymes, peroxiredoxin 3 (Prx3) and manganese-dependent superoxide dismutase (MnSOD) [22], which are transcriptional targets of FOXO3 [13] and Nrf2 [24]. However, it must yet be established which transcription factor/s mediate these effects.

Therefore, aiming to fully characterize the molecular mechanisms involved in the cardiovascular effects of (PhSe)_2_ and considering the relevance of metabolic alterations in cardiovascular diseases, we analyzed how (PhSe)_2_ impacted redox-bioenergetic cellular pathways under glycolytic or oxidative *in vitro* conditions in primary bovine vascular endothelial cells (BAEC).

## 2. Materials and Methods

### 2.1. Cell culture and treatments

BAEC were extracted from a fresh bovine thoracic aorta, obtained in an authorized slaughterhouse, as previously described by Peluffo and colleagues [25]. BAEC were cultured in growth medium (DMEM) supplemented with 10% of fetal bovine serum (FBS; Gibco/Invitrogen), containing 2 mM glutamine, 100 units/mL penicillin, 100 μg/mL streptomycin, 10 mM HEPES, 25 mM glucose, 44 mM NaHCO_3_ and incubated at 37°C in a humidified atmosphere with 5% CO_2_. Cell suspensions were seeded in 96, 24 or 6-well plates (100 × 20 mm), at different cell densities, depending on the experimental procedure. Cells at passages P4–P8 were used to ensure they maintained sufficient mitochondrial plasticity. Cells were allowed to reach 90% confluence and under *glycolytic or oxidative conditions*, as previously described [15], treated with 1 μM (PhSe)_2_ or vehicle (DMSO 0.05%) for 3, 6, 12, 24 or 48h. In short, when BAEC were submitted to the *glycolytic protocol*, culture media was changed to DMEM without FBS and treated with 1 μM (PhSe)_2_ or vehicle (DMSO 0.05%) for 3, 6, 12, 24 or 48h. When BAEC were submitted to the *oxidative protocol*, culture media was changed to glucose-free RPMI medium with 5 mM galactose and without FBS. Cells were maintained in this medium for 16 hours and then treated with 1 μM (PhSe)_2_ or vehicle (DMSO 0.05%) for 3, 6, 12, 24 or 48h. For each independent experiment, identical cells, derived from the same original culture dish, were split and then exposed to glucose and galactose media separately. Therefore, all the cells within each independent experiment had the same passage. The use of 1 μM (PhSe)_2_ dosage is based on previous studies from our group and others that demonstrated the protective effect of (PhSe)_2_ on BAECs at a concentration of 1 μM, as well as the absence of cytotoxic effects [16,17].

### 2.2. High-resolution respirometry by Seahorse

To evaluate mitochondrial oxygen consumption in BAEC, cells were plated in XF24 cell culture microplates (24-well plates at a cell density of 5 x 10^3^ cells/well), submitted to the glycolytic (glucose) or oxidative (galactose) protocols and then treated with 1 μM (PhSe)_2_ or vehicle for 3, 12 or 48h. A calibration cartridge (Seahorse Bioscience) was equilibrated overnight and then loaded with unbuffered DMEM (port A), 0.6 μM oligomycin (port B), 0.3 μM carbonyl cyanide 4- (trifluoromethoxy)phenylhydrazone (FCCP) (port C), and 0.1 μM rotenone plus 0.1 μM antimycin A (port D), all from Sigma-Aldrich. This allowed the determination of basal respiration, maximal respiration, reserve capacity, and non-mitochondrial respiration. In all experiments, the protein concentration in each well was determined at the end of the measurements, using the Pierce BCA protein assay kit (Thermo Scientific) following cell lysis with RIPA buffer (Sigma-Aldrich) supplemented with a protease inhibitor cocktail (Complete Mini; Roche), and used to calibrate the oxygen consumption data.

### 2.3. MitoSOX Imaging

Mitochondrial superoxide (O_2_^-^) was evaluated by MitoSOX Red (Molecular Probes, Carlsbad, CA, USA) staining, essentially as described [26]. Briefly, BAEC were grown on coverslips in 24-well culture (1 x 10^5^ cells/well) plates, submitted to the glycolytic (glucose) or oxidative (galactose) protocols and treated with 1 μM (PhSe)_2_ or vehicle (DMSO 0.05%) for 24h. Then, BAEC were exposed to 500 μM hydrogen peroxide (H_2_O_2_) for 4h or 80 μM 1,4-naphthoquinone (DMNQ) for 2h, and finally incubated with 3 μM MitoSOX Red for 10 min and fixed with paraformaldehyde (2%) and analyzed by fluorescence microscopy (Nikon 90i). Under our experimental conditions, no nuclear fluorescence was detected, indicating that the probe remained confined to the mitochondria. Therefore, nuclear counterstaining with DAPI or equivalent dyes was not required for image segmentation. For each slide, four images were acquired, and the total fluorescence intensity was determined using ImageJ software. The average fluorescence from the four images was then calculated and considered as the representative fluorescence value for one replica sample.

### 2.4. Immunofluorescence (IF)

BAEC were grown on coverslips, 24-well culture (1 x 10^5^ cells/well) plates and submitted to the glycolytic (glucose) or oxidative (galactose) protocols and then treated with 1 μM (PhSe)_2_ or vehicle for 3, 6, 12, 24 or 48h. At the end of the incubation period, cells were then fixed with 3.7 % formaldehyde, permeabilized with 0.1 % Triton, and incubated consecutively with a primary antibody Tomm22 (1:200, Sigma Life Science, cat: HPA003037), Nrf2 (1:200, Thermo Fisher, cat: PA5-27882) or FOXO3 (1:100, Cell signaling, cat: 9467) and then a secondary antibody (IgG rabbit ALEXA-488 conjugate, 1:2500). Finally, cells were counterstained with DAPI, mounted and examined under a confocal microscope (Zeiss LSM 700, Obercochen, Germany), as previously described [26].

Tomm22 was used as a marker to assess mitochondrial content and subcellular distribution using Image J software analysis of confocal microscopy images. Total Tomm22 positive area was considered to indicate mitochondrial volume. Subcellular distribution was analyzed by drawing a ling along the longitudinal cell axis and by calculating the ratio between the perinuclear Tomm22 signal (15 pixels from the nuclear edge) and the total cytosolic signal average provided by that line. Signal asymmetry in the perinuclear region was assessed by calculating the rate of the maximum and minimum intensity values on opposite sides of the nucleus as divided by the above-mentioned cross-section line. Using this line, mitochondrial fission was estimated by the calculation of the standard deviation of Tomm22 signal across the cytosol. The nucleus was identified using the DAPI staining signal, this analysis procedure was as previously described [15,27]. Since BAECs are primary cells with significant variability in mitochondrial and respiratory parameters depending on passage number [15], all data were normalized to the t = 0 values of each replica and expressed as a percentage in the graphs.

For the analysis of nuclear and cytosolic content of Nrf2 and FOXO3, as previously described [15], a similar procedure was used, using Imagen J Software, a line was drawn across the long axis of each cell the cellular compartment and the nuclear region was identified based on DAPI fluorescence signal. The Nrf2 average fluorescence signal corresponding to the and cytosolic regions was then calculated and then ratio of both, resulting in the nuclear-cytosolic ratio.

### 2.5. Protein extraction and Western blotting

BAEC (5 x 10^5^ cells/well) were grown in 6-well culture plates, submitted to the glycolytic (glucose) or oxidative (galactose) protocols and treated with 1 μM (PhSe)_2_ or vehicle for 3, 6, 12, 24 or 48h. The cells were then washed with phosphate-buffered saline (PBS) and lysed in 150 μl lysis buffer containing 150 mM NaCl, 0.1% sodium dodecyl sulphate (SDS), 1% sodium deoxycholate, 1% NP40 and 25 mM Tris–HCl pH 7.6, in the presence of protease (Complete, Roche Diagnostics, Mannheim, Germany) and phosphatase inhibitors (Sigma-Aldrich, St. Louis, MO). Cells were harvested by scraping, samples were clarified by centrifugation at 13.000 rpm for 15 min at 4 °C and protein concentration was determined using the BCA assay (Thermo Scientific, Rockford, IL, USA). Equal amounts of protein (40 – 50 μg) from whole extracts were separated by SDS-PAGE electrophoresis, using 10/12% acrylamide/bisacrylamide (29:1) 0.1% SDS gels and transferred to a 0.2 μm PVDF Hybond-P membranes (Amersham/GE healthcare) at 20V for 1h in a semi-dry Trans-Blot SD cell system (Bio-Rad, Hercules, CA). Following the blockade of the membranes using TBS-T (20 mM Tris, 150mM NaCl, 0.1% Tween 20) with 3% BSA, they were incubated with primary antibodies and then secondary antibodies in blocking solution, FOXO3 (1:1000, Cell signaling, cat: 9467), pFOXO3 (1:1000, Cell signaling, cat: 2309), Akt (1:1000, Cell signaling, cat: 2920), pAkt (1:1000, Cell signaling, cat: 4060), MnSOD (1:3000, ADI-SOD-110, Enzo Life Sciences, cat: ADI-SOD-110-F), Prx3 (1:1000; LabFrontier, Daehyun-dong Seodaemun-gu Seoul, Korea, cat: LF-PA0030), GCLC (1:1000; Cell Signaling, cat: 48005), GCLM (1:1000; Sigma-Aldrich, cat: SAB5700737) and β-Tubulin (1:80000; Sigma, St. Louis, M, cat: T9026), that was used as loading control. Secondary antibodies rabbit and mouse IgG were from LI-COR Biosciences (Lincoln, NE). Membranes were then extensively washed with TBS-T and developed using Bio-Rad Clarity Western ECL Substrate. Images were captured using a luminometer scanner and subject to densitometry analysis.

### 2.6. Image analysis

Image J software was used for the analysis of areas, signals in an area and cross section signals from fluorescence and confocal microscopic images for Tomm22, Nrf2, FOXO3 and MitoSOX and to analyze Western blotting membranes.

### 2.7. Gene expression analysis of genes coding for antioxidant proteins by RT-qPCR

To evaluate mRNA levels, BAEC (5 x 10^5^ cells/well) were seeded in 6-well plates and cultivated for 24h. Following cell treatment, total RNA was isolated using TRIzol Reagent (Invitrogen, Carlsbad, CA) following the manufacturer instructions. 1 μg of total RNA was then reverse transcribed by extension of random primers using M-MLV (Promega, Madison,WI). Relative expression levels were determined by real-time PCR (qPCR) in a 7900 Sequence Detection System (Applied Biosystems, Carlsbad, CA) using the primer sets listed below/in Table 1.

**Table 1.**
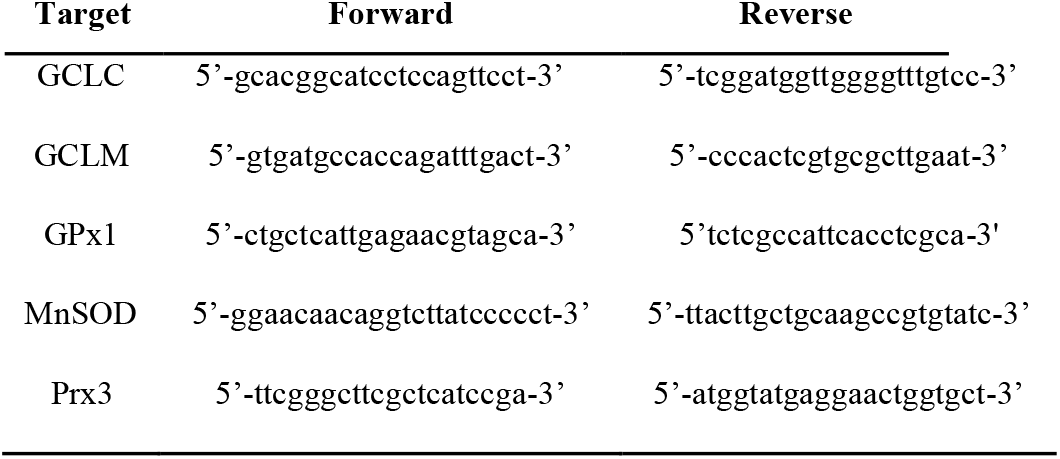
Primer pairs used in RT-qPCR to quantify gene expression changes.

### 2.8. Statistical analysis and graphics

Numerical raw data was compiled and processed in Excel files (Microsoft) and then final data was incorporated into GraphPad PRISM® software version 6.0 for Windows (GraphPad Software, San Diego, CA, USA), that was used for statistical analysis and preparation of graphs. Normal (Gaussian) distribution was evaluated with the Shapiro-Wilk normality test. Significance of differences among data sets was evaluated using unpaired test *t* or two-way of variance analysis (ANOVA), using Bonferroni’s correction for multiple comparisons. Results are expressed as mean ± SD. *P* <0.05 was considered statistically significant. n ≥ 3 in all experiments. The number of independent experiments is indicated in the corresponding figure legends. Each data point incorporated in the graphs represents an independent experiment, not technical replicas. Independent means they used an independent vial of cells, thaw on a different day and thus tested on different days. Data from technical replicas/individual cells/individual images were always pooled to calculate each data point. The graphical abstract was created in BioRender De bem, L. (2025) https://BioRender.com/pu1070I under the agreement number XK28BVWXKS.

## 3. Results

### 3.1. Effect of (PhSe)_2_ on mitochondrial respiration in BAEC

We first tested the differential effect of (PhSe)_2_ on mitochondrial respiration in BAEC cultured in standard glucose-containing media or in galactose-containing medium to force the cells to rely on OXPHOS for adenosine triphosphate (ATP) production. All control data comparing alterations in BAECs cultured in glucose-versus galactose-containing media have been recently published in [15]. In that study we observed that the maximum oxidative capacity after 12–48 h of incubation of BAEC conditioned in culture medium containing galactose was significantly higher when compared to that observed in cells in culture medium containing glucose. Now, we found that 3h treatment with (PhSe)_2_ decreased maximum respiration in both glucose- (*F* (1, 19) = 4,959, *p* = 0.0382) and galactose- (*F* (2, 12) = 10,56, *p* = 0.0023) containing medium (Fig. 1A, C and D) but the oxygen consumption rate (OCR) recovered after 48h of treatment with (PhSe)_2_ in both glucose- and galactose-containing medium (Fig. 1B, C and D). Furthermore, when the maximum OCR was normalized for the mitochondrial volume, estimated by immunofluorescence analysis of the mitochondrial protein Tomm22, we found that this ratio was significantly increased following 48h of treatment with (PhSe)_2_ in glucose-containing medium (*t* = 3.512, *p* = 0.0246), suggesting a higher respiratory capacity per mitochondria in this context (Fig. 1E). However, in a galactose-containing medium the treatment with (PhSe)_2_ did not significantly change the maximum respiration/mitochondrial volume unit ratio (Fig. 1F).

**Fig. 1.**
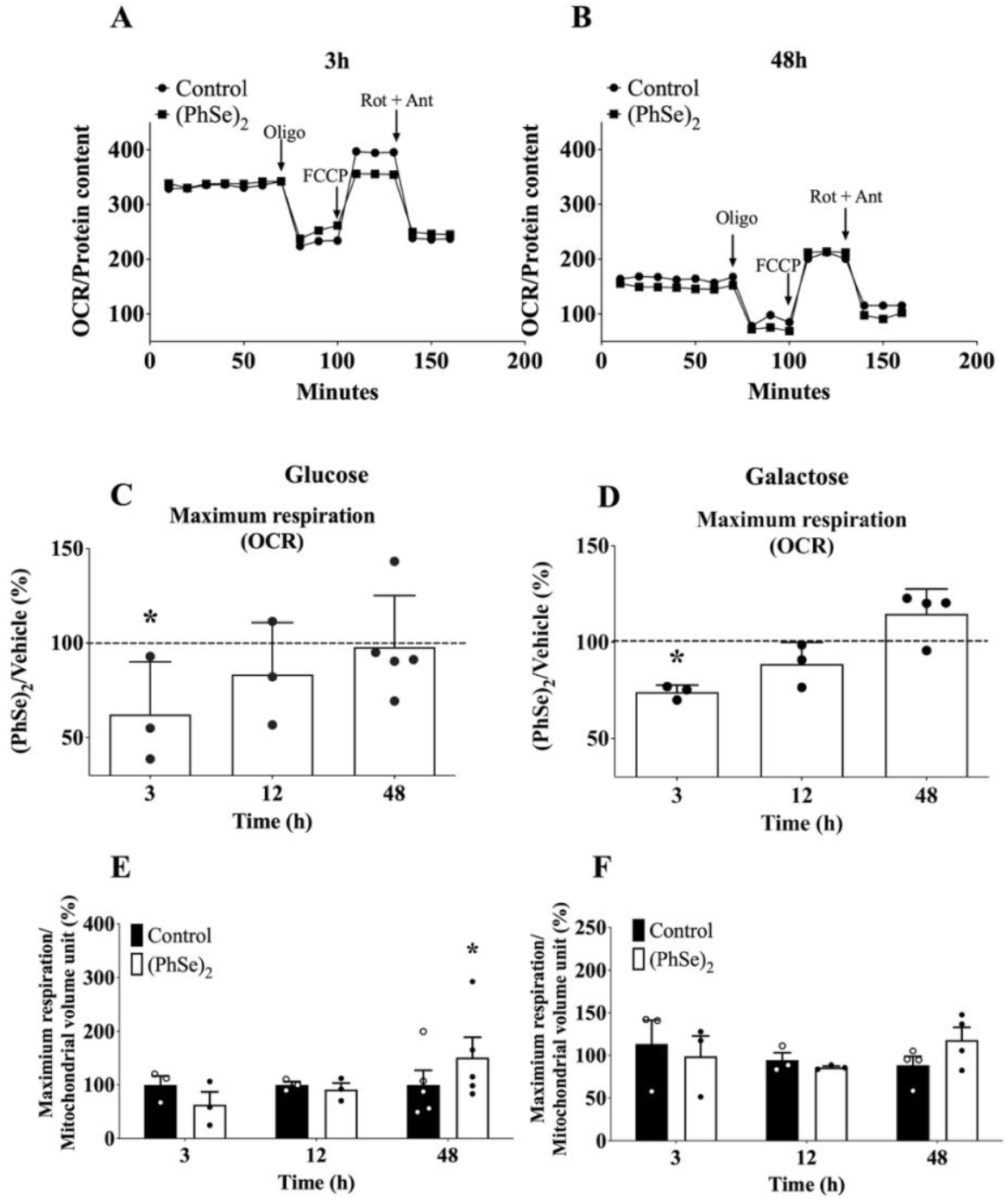
Effect of (PhSe)_2_ on mitochondrial respiration in BAEC conditioned in glucose- or galactose-containing medium. (A) Representative respirometry assay of BAEC conditioned in glucose-containing medium for 3h and (B) 48h, data corrected for protein amount. (C) Maximum respiration in BAEC conditioned in glucose-containing medium. (D) Maximum respiration in BAEC conditioned in a galactose-containing medium. (E) Ratio of maximum respiration per mitochondrial volume unit in BAEC treated with (PhSe)_2_ for 3, 12 or 48h in glucose-containing medium and (F) galactose-containing medium. (A and B) Plots of O_2_ consumption (OCR). (C and D) Quantification was corrected using total μg of protein and expressed as the percentage of treatment with (PhSe)_2_ (white bars) with respect to vehicle group (dashed line). Average points recorded in samples with (PhSe)_2_ after the addition of FCCP/average points recorded in samples with vehicle after the addition of FCCP x100. (E and F) The mitochondrial volume unit was analyzed by Tomm22 immunofluorescence in 5 independent experiments (Figure 2). The average of the samples after FCCP by Seahorse/average of Tomm22 was analyzed by immunofluorescence. Data were represented as mean ± SD (n=3-5). (C and D) \**p* < 0.05, indicate statistical difference from the vehicle group by two-way ANOVA, followed by Bonferroni’s *post hoc* test, (E and F) \**p* < 0.05, indicates statistical difference from vehicle group by paired t-test.

### 3.2. Effect of (PhSe)_2_ on mitochondrial dynamic in BAEC conditioned in glucose or galactose medium

To understand the changes elicited in the mitochondria by (PhSe)_2,_ we monitored mitochondrial dynamics by immunofluorescence analysis of Tomm22 using confocal microscopy. Total mitochondrial volume, fusion, and subcellular distribution were analyzed in BAEC treated with (PhSe)_2_ for 3, 12, and 48h in glucose- or galactose-containing medium. We found that in glucose-containing medium (Fig 2A-E), although the treatment with (PhSe)_2_ for 48h decreased total mitochondrial content (*t* = 3.161, *p* = 0.0195) (Fig. 2B), it increased mitochondrial fusion (*t* = 3.022, *p* = 0.0233) (Fig. 2C), reduced the mitochondrial content in the perinuclear region (*t* = 3.218, *p* = 0.0182) (Fig. 2D), consistent with the observed increase in oxidative capacity per mitochondria. Furthermore, it increased the asymmetrical distribution of mitochondria within the cells, which could be related to intercellular coupling/organization of the epithelium polarity (*t* = 3.506, *p* = 0.0172) (Fig. 2E). In contrast, cells maintained in galactose-containing medium (Fig. 2F-J) did not change the total mitochondrial content (Fig. 2G), fission (Fig. 2H), or mitochondrial localization in the perinuclear region (Fig. 2I) in response to the treatment with (PhSe)_2_. However, a transient increase in the asymmetrical distribution of mitochondria was observed at 12h (*t* = 3.041, *p* = 0.0287), which reverted after 48h (*t* = 5.916, *p* = 0.001) of treatment (Fig. 2J).

**Fig. 2.**
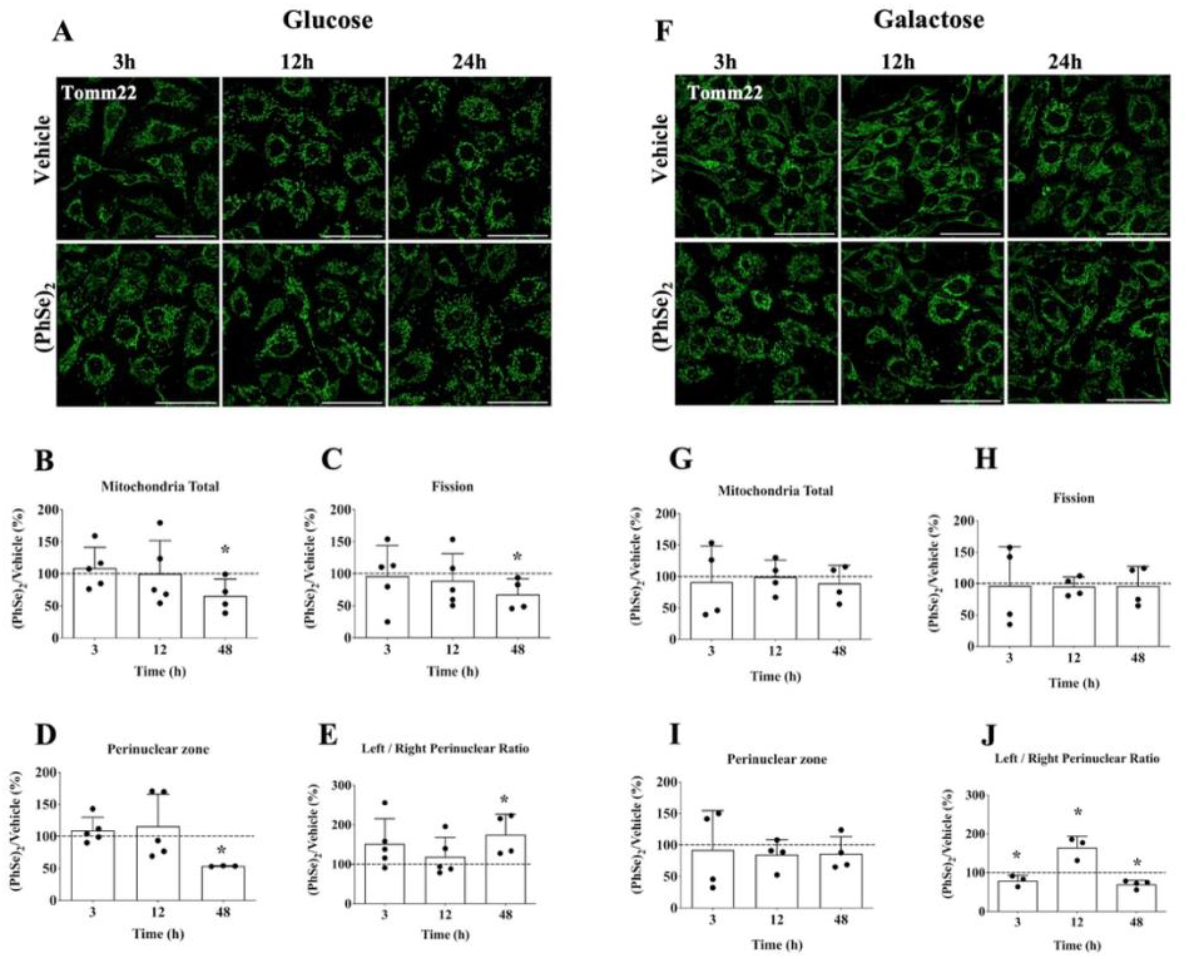
Effect of (PhSe)_2_ on mitochondrial dynamics in BAEC conditioned in glucose- or galactose-containing medium. BAEC were treated with (PhSe)_2_ (1 μM) or vehicle (DMSO) for 3, 12, or 48h in medium containing glucose (A-E) or galactose (F-J), and the mitochondrial parameters were determined by immunofluorescence of Tomm22 antibody. (A and F) Representative image of mitochondrial Tomm22 (green) in BAEC treated with (PhSe)_2_. The white bars represent 50 μm. (B and G) Quantification of mitochondrial volume, (C and H) mitochondrial fission, (D and I) intensity of the perinuclear zone, (E and J) ratio of left/right intensity. Results were expressed as the percentage of treatment with (PhSe)_2_ (white bars) with respect to the vehicle group (dashed line). Data were represented as mean ± SD (n=3-4). \**p* < 0.05, indicates statistical difference from vehicle group by unpaired t-test.

### 3.3. (PhSe)_2_ prevents the mitochondrial superoxide production in BAEC exposed to oxidants

To evaluate if (PhSe)_2_ could modify the cellular capacity to respond to an oxidative challenge, we pre-treated BAEC with 1 μM of (PhSe)_2_ and then exposed the cells to H_2_O_2_ (500 μM) or DMNQ (80 μM), an inhibitor of the mitochondrial electron transport chain that drives the formation of mitochondrial superoxide (O_2_^-^). Mitochondrial RS generation was evaluated using the MitoSOX probe. In BAEC cultured in glucose-containing medium, DMNQ significantly increased O_2_^-^ production. Importantly, (PhSe)_2_ pre-treatment reduced mitochondrial O_2_^-^ levels in DMNQ challenged cells (*F* (1,8) = 7.35, *p* = 0.0266). H_2_O_2_ in glucose-cultured cells also increased mitochondrial O_2_^-^ levels and, consistently, (PhSe)_2_ pre-treatment reduced H_2_O_2_ levels (*F* (1,7) = 13.66, *p* = 0.0077) (Fig. 3A-B), suggesting that (PhSe)_2_ pre-treatment improved mitochondrial homeostasis in glucose cultured BAEC. In contrast, in BAEC cultured in galactose-containing medium, DMNQ did not generate a significant increase in O_2_^-^ production, possibly because in these conditions the mitochondria are more resistant to DMNQ action, while H_2_O_2_ did. Furthermore, pre-treatment with (PhSe)_2_ reduced mitochondrial O_2_^-^ levels following H_2_O_2_ exposure (*F* (1,7) = 9.171, *p* = 0.0192) (Fig. 3C-D). These results consistently support the idea that the (PhSe)_2_ reduces the mitochondrial accumulation of O_2_^-^ levels.

**Fig. 3.**
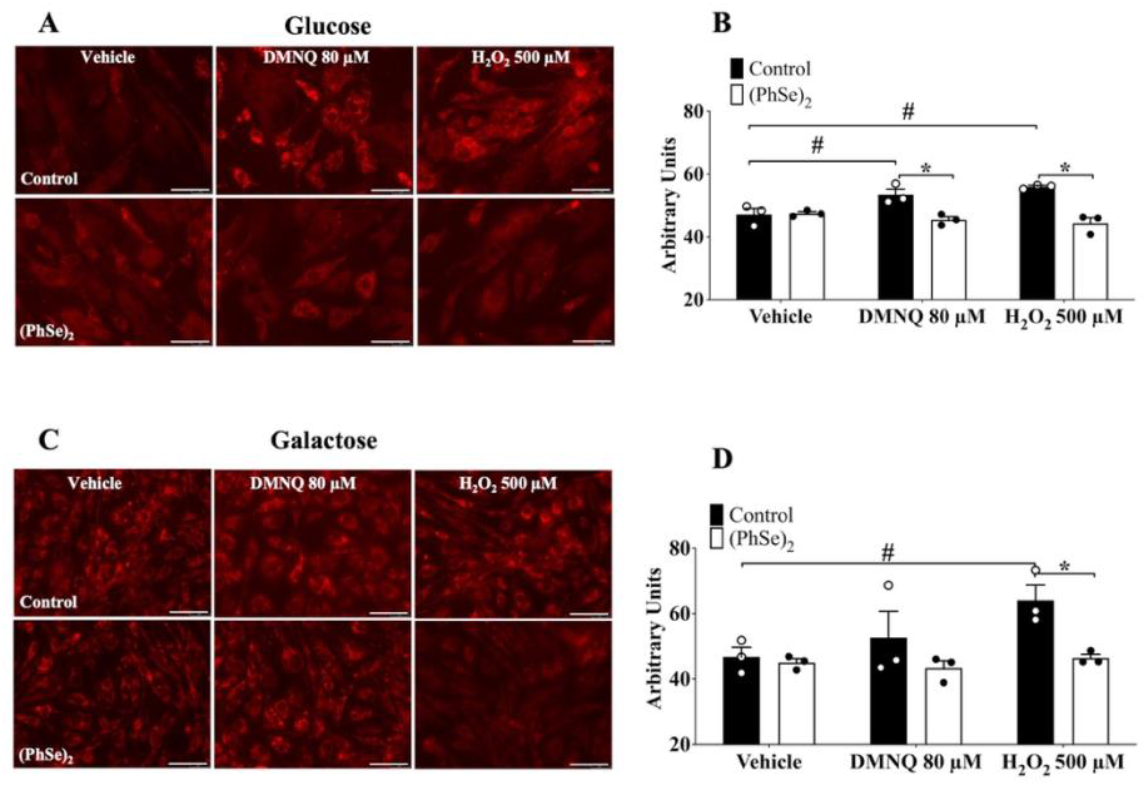
Effect of (PhSe)_2_ on mitochondrial O_2_^-^ production in BAEC conditioned in glucose- or galactose-containing medium. BAEC were pretreated with (PhSe)_2_ (1 μM) or vehicle (DMSO) for 24h in medium containing glucose (A-B) or galactose (C-D) and subjected to oxidants as DMNQ or H_2_O_2_. The O_2_^-^ production was determined by Nikon 90i microscopy through the MitoSOX probe. (A and C) Representative image of mitochondrial O_2_^-^ production in BAEC pretreated with (PhSe)_2_ and exposed to DMNQ (80 μM) or H_2_O_2_ (500 μM) (for 2 or 4h, respectively), the white bars represent 50 μm. (B and D) Quantification of mitochondrial O_2_^-^ production (values x 10^4^) in BAEC pretreated with (PhSe)_2_ and exposed to DMNQ or H_2_O_2_. * *p* < 0.05, indicates statistical difference between the groups exposed to oxidants and (PhSe)_2_, # *p* < 0.05, indicates the statistical difference between vehicle and DMNQ or H_2_O_2_ groups by two-way ANOVA, followed by Bonferroni’s *post hoc* test.

### 3.4. Time-course effect of (PhSe)_2_ in the expression of antioxidant proteins in BAEC conditioned in glucose or galactose medium

Next, we decided to evaluate if the observed effect of (PhSe)_2_ on mitochondrial O_2_^-^ levels could be derived from an increased cellular detoxification capacity. To that end, we used targeted gene expression and protein analysis of some major antioxidant coding genes regulated by the two crucial redox-sensitive transcription factors, Nrf2 and FOXO3. In BAEC cultured in glucose-containing medium, following 6h of treatment with (PhSe)_2_ we observed a significant increase in MnSOD and *SOD2* gene expression levels (*F* (3,47) = 5.159, *p* = 0.0036 and *F* (1,29) = 7.157, *p* = 0.0121, respectively) (Fig. 4A-B) as well as an increase in Prx3 protein levels (*F* (3,36) = 2.722, *p* = 0.053) (Fig. 4C-D). Both MnSOD and Prx3 are antioxidant proteins that localize in the mitochondrial matrix. We also monitored the effect of (PhSe)_2_ on GCL, the enzyme of the limiting step for the synthesis of the antioxidant tripeptide glutathione, and GPx1, the main enzyme involved in the recovery of reduced glutathione following its oxidation, in glucose-containing medium. GCL is made up of two subunits. We noted that GCLC protein and *GCLC* gene expression levels increased (*F* (1,20) = 30.42, *p*<0.0001 and *F* (1,27) = 16.94, *p* = 0.0003, respectively) following 24h of treatment with (PhSe)_2_ (Fig. 4E-F) as well as *GCLM* levels at 6h of treatment (*F* (1,30) = 8.759, *p* = 0.006) (Fig. 4 G-H). Furthermore, *GPx1* levels were increased at 12 and 24h of treatment with (PhSe)_2_ in BAEC cultured in glucose-containing medium (*F* (1,14) = 29.11, *p* < 0.0001) (Supplementary Fig. 1A). Similar results were observed in BAEC maintained in a galactose-containing medium.

**Fig. 4.**
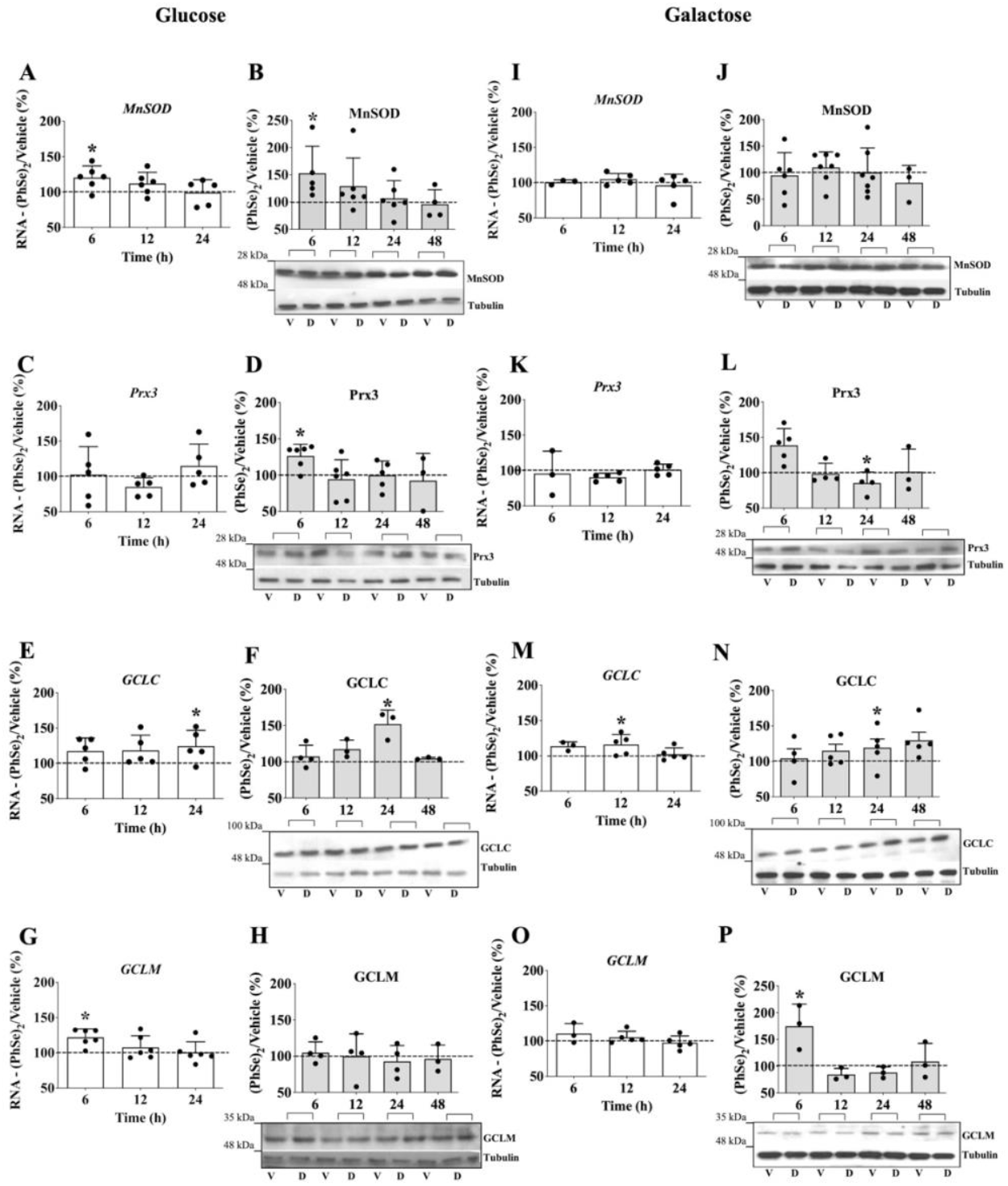
Effect of (PhSe)_2_ in time-course expression profiles of antioxidant proteins in glucose- or galactose-containing medium. BAEC were treated with (PhSe)_2_ or vehicle (DMSO) for 6, 12, 24 or 48h in glucose or galactose medium. Protein expression by Western Blotting analyses and mRNA expression by qPCR. Results were expressed as protein /β-tubulin ratio and the data were presented as the percentage of treatment with (PhSe)_2_ (bars) with respect to vehicle group (dashed line). V = vehicle, D = (PhSe)_2_. Data were represented as mean ± SD (n=3-7). \**p* < 0.05, indicate statistical difference from vehicle group by two-way ANOVA, followed by Bonferroni’s *post hoc* test.

We observed an increase in the mRNA levels of *GCLC* after 12h (*F* (1,22) = 14.07, *p* = 0.0011) and GCLC protein content at 24h (*F* (1,31) = 9.316, *p* = 0.0046) (Fig. 4 M-N), as well as an increase in GCLM protein levels after 6h of treatment with (PhSe)_2_ (*F* (3,14) = 5.9, *p* = 0.0076) (Fig. 4 O-P). (PhSe)_2_ also increases the mRNA expression of GPx1 at 12 and 24h of treatment in the galactose-containing medium (*F* (1,12) = 40.22, *p* < 0.0001) (Supplementary Fig. 1B). In contrast, *SOD2* mRNA levels and protein of MnSOD were unaltered (Fig. 4 I-J), while the Prx3 levels decreased after 24h of treatment with (PhSe)_2_ in the galactose-containing medium (Fig. 4 K-L). The mRNA levels of catalase (*CAT*), mitochondrial uncoupling protein 2 (*UCP2*), cytochrome c (*CYTC*), glutathione synthetase (*GSS)* and *Nrf2* were unaltered by the treatment with (PhSe)_2_ of BAEC maintained in either glucose- or galactose-containing medium (Supplementary Fig. 1 C-L). Overall, these results suggest that in glucose containing media (PhSe)_2_ induces both the main mitochondrial and cytosolic antioxidant systems, while in galactose media (PhSe)_2_ induces mainly the cytosolic antioxidant systems, a result consistent with the expected pre-activation of OXPHOS and mitochondrial antioxidant capacity in galactose media.

### 3.5. Effect of (PhSe)_2_ on Nrf2 nuclear translocation of BAEC conditioned in glucose or galactose medium

Since (PhSe)_2_ increased the levels of antioxidant genes known to be targeted by the transcription factor Nfr2, we evaluated the effect of (PhSe)_2_ on Nrf2 nuclear translocation, a thus activation, in BAEC cultured in glucose- or galactose-containing medium. We found that Nrf2 nuclear localization was increased following 3h of incubation with (PhSe)_2_ in BAEC cultured in both glucose and galactose containing medium (*F* (4,14) = 4.737, *p* = 0.0126 and *F* (1,30) = 64.98, *p* < 0.0001, respectively) (Fig. 5A-B). The increased nuclear localization was gradually reduced overtime in both glucose and galactose cultured cells but was maintained over a more extended period in galactose cultured cells (Fig. 5C-D), suggesting a more effective nuclear retention in galactose. The separate evaluation of nuclear and cytosolic levels of Nrf2 showed a significant increase in cytosolic Nrf2 levels following 3h of treatment with (PhSe)_2_ in BAEC maintained in glucose-containing medium but not in galactose cultured cells possibly suggesting that global Nrf2 levels stabilized by (PhSe)_2_ in glucose but not in galactose media (Supplementary Fig. 2). Nrf2 degradation is a tightly regulated process, cytosolic retention of Nrf2 by Keap1 is linked to its proteasomal degradation, and therefore, Keap1 release is generally associated not only to Nrf2 nuclear translocation but also to its stabilization [28]. In summary, Nrf2 activation is detectable in both glucose and galactose media, but in galactose media is likely to be effective for more extended periods of time.

**Fig. 5.**
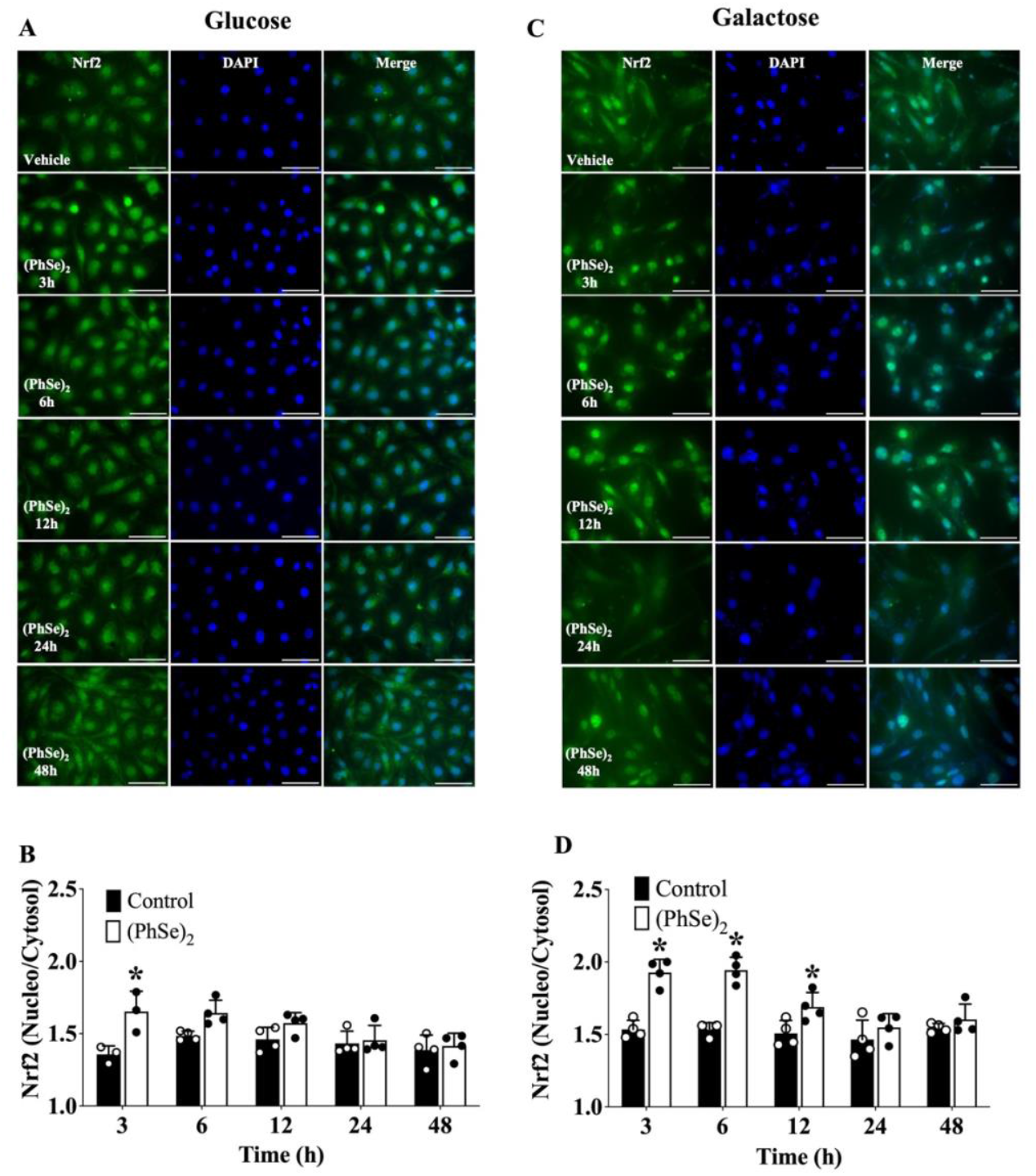
Effect of (PhSe)_2_ on Nrf2 translocation in BAEC conditioned in glucose- or galactose-containing medium. BAEC were treated with (PhSe)_2_ (1 μM) or vehicle (DMSO) for 3, 6, 12, 24 and 48h in medium containing glucose (A-B) or galactose (C-D) and the immunofluorescence the Nrf2 were analyzed. (A and C) Representative image of immunofluorescence from Nrf2 (green) and DAPI (blue) in BAEC, the white bars represent 50 μm. (B and D) Ratio of Nrf2 in the nuclei/cytosol. Data were represented as mean ± SD (n=3-4). * *p* < 0.05, indicate statistical difference from vehicle group by two-way ANOVA, followed by Bonferroni’s *post hoc* test.

### 3.6. Effect of (PhSe)_2_ on FOXO3 activation in BAEC conditioned in glucose or galactose medium

Finally, we also analyzed the possible involvement of FOXO3 transcriptional factor in (PhSe)_2_ activation of antioxidant gene expression. Treatment of BAEC in glucose-containing medium with (PhSe)_2_ for 6, 12 or 24h increased the level of FOXO3 phosphorylation (pFOXO3) at Thr32 (*F* (1,90) = 8.111, *p* = 0.0055), a mark of Akt phosphorylation (pAkt), suggesting an inhibition of FOXO3 activity and nuclear extrusion (Fig. 6A). However, no significant changes in the nuclear/cytosolic ratio of FOXO3 (Fig 6E) nor in the cytosolic and nuclear levels when analyzed separately (Supplementary Figure 3), and, although an increase in Akt phosphorylation was detectable in response to (PhSe)_2,_ it could only be observed at 24h of treatment in glucose media (*F* (1,76) = 12.86, *p* = 0.0006) (Fig. 6C). In contrast, although the nuclear/cytosolic ratio of FOXO3 transiently decreased at 3h of treatment in galactose media (Fig. 6F), (PhSe)_2_ decreased pFOXO3/FOXO3 ratio at 24h (*F* (3,57) = 5.642, *p* = 0.0019), suggesting an increase in FOXO3 activity and nuclear localization (Fig. 6B). In these conditions although the changes in pAkt levels did not reach statistical significance, a consistent tendency to lower pAkt/Akt ratio was detectable at 24h (Fig. 6D) as well as detectably higher FOXO3 levels in both the nuclei and the cytosol at 12h of treatment (Supplementary Figure 3). All together these results suggest that FOXO3 activity is more likely to play a relevant role in antioxidant gene regulation in galactose than in glucose cultured cells.

**Fig. 6.**
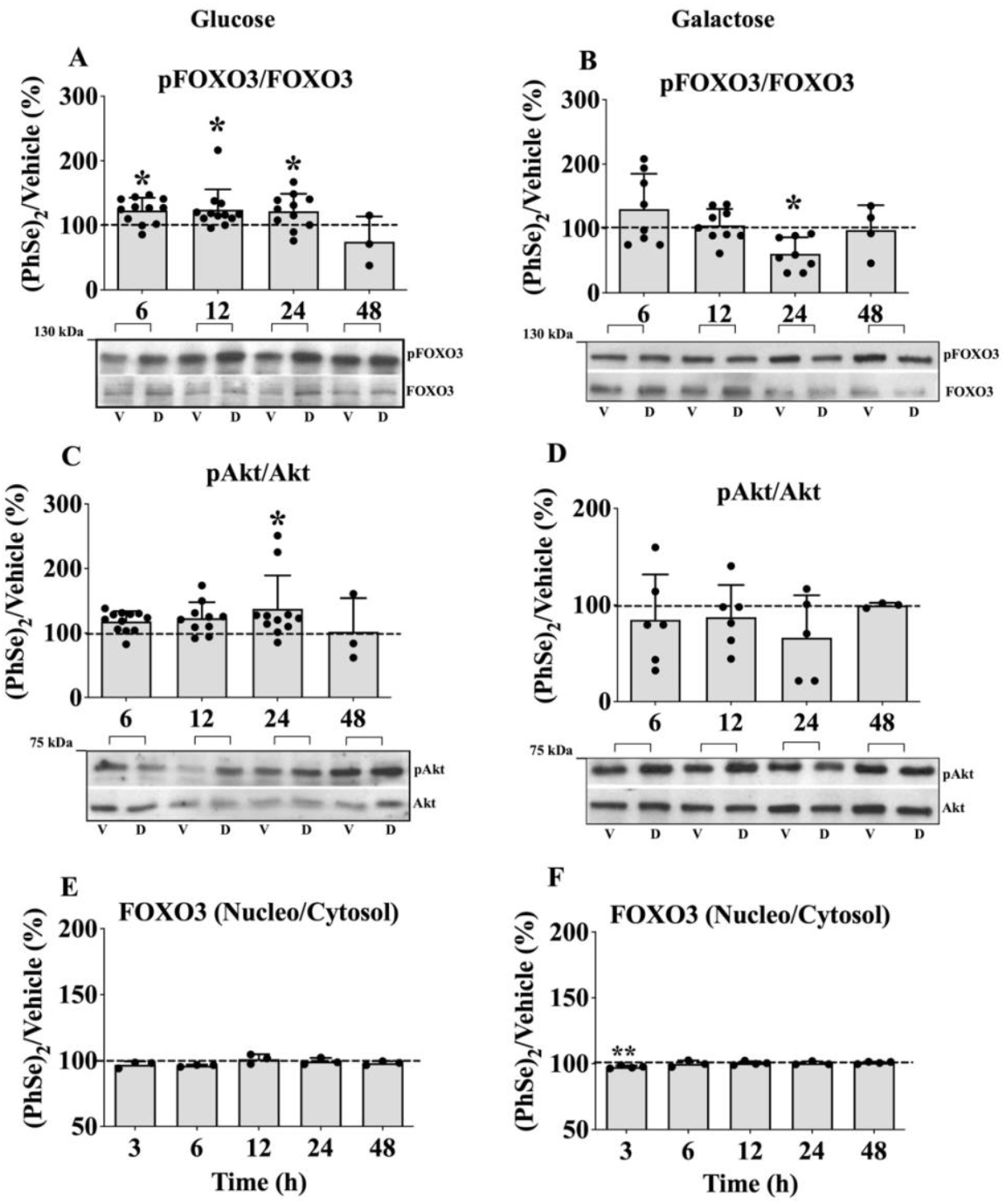
Effect of (PhSe)_2_ on pFOXO3 and pAkt levels in BAEC conditioned in glucose- or galactose-containing medium. BAEC were treated with (PhSe)_2_ or vehicle (DMSO) for 3, 6, 12, 24 and 48h in glucose (A, C and E) or galactose (B, D and F) medium to analyze pFOXO3 (A-B) and pAkt (C-D) protein content by Western blotting. Results are expressed as protein/β-tubulin or FOXO3 ratio and the data is presented as the relative % of treatment with (PhSe)_2_ (bars) *vs* vehicle. V = vehicle, D = (PhSe)_2_. (E and F) Immunofluorescence for FOXO3 were analyzed. Ratio of FOXO3 in the nuclei/cytosol. Data were represented as mean ± SD (n=3-13). * *p* < 0.05, indicate statistical difference from vehicle group (dashed line) by two-way ANOVA, followed by Bonferroni’s *post hoc* test.

## 4. Discussion

Endothelial cells are the primary target of circulating redox-active molecules. Among these, primary bovine aortic endothelial cells (BAEC), since preserving their original metabolic plasticity, represent a relevant *in vitro* model for investigating the effects of pharmacological agents across diverse metabolic contexts. The choice of cell culture media profoundly influences the metabolic performance of cells *in vitro*. Our recent findings, particularly concerning the modulation of transcription factors Nrf2 and FOXO3 key regulators of antioxidant gene expression in BAECs, underscore the significance of cellular metabolic state. Specifically, alterations in cell culture conditions, leading to heightened reliance on oxidative metabolism, demonstrate a consequential impact on the efficacy of redox-active agents on vascular cells. These observations suggest that the effectiveness of redox-active agents on endothelial cells may be intricately linked to their metabolic milieu. Therefore, we investigate the effects of (PhSe)_2_ on BAEC cultured in media containing glucose or galactose, assessing its impact on metabolism and antioxidant responses.

(PhSe)_2_ has been extensively studied for its pivotal role in vascular physiology and pathology, attributed to its capacity for redox modulation. This includes the promotion of mitochondrial respiration and the upregulation of antioxidant gene expression, partly mediated by the translocation of Nrf2 to the nucleus [22]. Our findings reveal significant differences in the action of (PhSe)_2_ depending on whether BAEC are cultured in galactose and glucose media. These differences are likely to be relevant to their vascular effects *in vivo*. In cells cultured in glucose, treatment with (PhSe)_2_ yielded a net positive impact on mitochondrial oxidative capacity, assessed by respirometry. This enhancement correlated with observed changes in mitochondrial cellular dynamics, characterized by more fused mitochondria and a uniform distribution of mitochondria throughout the cells, with reduced concentration in the perinuclear region. Conversely, these changes were not observed in cells cultured with galactose, where the cells were already compelled to utilize their oxidative capacity. Therefore, the modulation of mitochondrial function by (PhSe)_2_ appears to be effective only in cells exhibiting limited utilization of their mitochondrial oxidative capacity. Regarding (PhSe)_2_ antioxidant capacity, we noted that although (PhSe)_2_ induces the nuclear translocation of Nrf2 in both glucose and galactose media, Nrf2 remains longer in the nuclei of cells cultured in galactose media. This observation suggests a potential preference for nuclear retention in the galactose environment. However, any speculation regarding the underlying mechanism at this stage would be highly conjectural. Potential explanations could involve interactions with other transcription factors [29] or the inhibition of its nuclear extrusion [30].

The enhanced Nrf2 activation profile observed in galactose media is even more pronounced in the case of FOXO3a, where its activation is solely detectable in galactose media, with reduced levels of pFOXO3a and increased nuclear protein levels. This shared pattern suggests a more efficient induction of antioxidant gene expression in galactose compared to glucose media. However, analysis of the gene expression changes elicited by (PhSe)_2_ on crucial antioxidant genes controlling mitochondrial antioxidant capacity and glutathione synthesis and recycling reveals a stronger induction in glucose media. In galactose media, the changes induced by (PhSe)_2_ are not only smaller but also predominantly focused on glutathione homeostasis, suggesting a more cytosolic, rather than mitochondrial effect. Therefore, the induction of antioxidants genes by (PhSe)_2_ in glucose media likely requires additional transcription factors that are especially active in conditions of low mitochondrial activity. Potential candidates include NFkB [31] a redox-sensitive transcription factor controlling antioxidant gene expression, and c-MYC [32] among others. Additionally, since (PhSe)_2_ also enhances mitochondrial fission and maximal oxidative capacity, other pathways may be involved. Acute inhibition of mitochondrial respiration could drive compensatory mechanisms, such as the activation of PGC-1α, which can induce both antioxidant gene expression and mitochondrial fusion [8] while supporting FOXO3 [13] and Nrf2 [14] activation.

Despite the apparent lower capacity of (PhSe)_2_ to induce antioxidant gene expression in galactose media, assessments of mitochondrial antioxidant capacity using the mitochondrial inhibitor DMNQ or the exogenous addition of H_2_O_2_ suggest a potentially heightened antioxidant response in galactose-treated cells. This inference arises from the substantial reduction elicited by (PhSe)_2_ on mitochondrial superoxide (O_2_^-^) levels in galactose-cultured cells following a challenge with H_2_O_2_. However, drawing definitive conclusions from these observations is challenging. In galactose media, cells exhibit greater resistance to DMNQ, while in glucose media, they are highly susceptible to H_2_O_2_-induced mitochondrial damage, resulting in under basal levels of mitochondrial O_2_^-^. Notably, (PhSe)_2_ partially rescues these aberrant levels and restores them to normal, consistent with its observed capacity to enhance mitochondrial oxidative capacity.

Therefore, our data suggest that while (PhSe)_2_ demonstrates greater efficacy in the activation of Nrf2 and FOXO3a under pro-oxidative conditions, alternative pathways contribute to its enhancement of mitochondrial oxidative capacity and total antioxidant capacity when cells are cultured in glucose media. The observation that (PhSe)_2_ induces acute inhibition of mitochondrial respiration shortly after treatment may indicate a significant role for mitochondrial function in this context. In summary, our findings underscore the critical influence of the cellular metabolic status on antioxidant capacity of redox-active molecules such as (PhSe)_2._

## Acknowledgments

This work was supported by MCIN/AEI/10.13039/501100011033/FEDER, UE, [grant numbers RTI2018-093864-B-I00 and PID2021-122765OB-I00]; the European Union’s Horizon 2020 research and innovation programme under the Marie Skłodowska-Curie [grant agreement 721236-TREATMENT]; EU Horizon Europe Program under the EIC Pathfinder DiBaN [GA-101162517]; the Brazilian institutions: Conselho Nacional de Desenvolvimento Científico e Tecnológico (CNPq), Coordenação de Aperfeiçoamento de Pessoal de Nível Superior (CAPES), Fundação de Apoio à Pesquisa do Distrito Federal (FAPDF) [grants 00193-00000884/2021-89 and 00193-00002348/2022-07]; and Instituto Nacional de Ciência e Tecnologia e Neuro-ImunoModulação (INCT-NIM) [grant 485489/2014-1]. The authors are grateful to Smart Servier Biology Medicine for the available figures.

